# Spatial lipidomics suggests fatty acid scavenging by an intranuclear parasite

**DOI:** 10.64898/2026.09.06.749766

**Authors:** Malin Stüwe, Benedikt Geier, Dolma Michellod, Miguel Ángel González Porras, Nicole Dubilier, Manuel Liebeke

## Abstract

Bacteria of the family *Endozoicomonadaceae* are widely associated with marine animals, yet their *in situ* metabolism remains mostly unknown. Genomic studies suggest that members of this family engage in extensive metabolic exchange with their hosts, ranging from nutrient provision in mutualistic sponge and coral symbioses to strong dependence on host metabolites like lipids in parasitic associations. Here, we study the lipid phenotype of the parasitic bacterium *Candidatus* Endonucleibacter childressi in its native tissue environment. The bacterium infects epithelial cells in the gill of the deep-sea mussel *Gigantidas childressi* and co-occurs with the mussel’s chemosynthetic symbionts providing a system to compare lipid profiles of symbiotic and parasitic gammaproteobacteria. We linked bacterial identity to metabolic phenotypes by combining mass spectrometry imaging with 16S rRNA fluorescence in situ hybridization. We identified phospholipids characteristic of regions infected with *Ca*. Endonucleibacter childressi. A subset of these lipids was also detected in symbiont-colonized regions, revealing a partial overlap in the phospholipid profiles of the two gammaproteobacteria. Fatty acids from PEs assigned to *Ca*. Endonucleibacter childressi contained polyunsaturated fatty acids such as FA 16:3, FA 18:3 or FA 18:4 which have not been reported from cultured close relatives. We hypothesize that the observed fatty acid profile may reflect adaptations to a host-associated lifestyle which involves the acquisition and remodeling of host-derived fatty acids. Our findings demonstrate the value of characterizing bacterial metabolic phenotypes directly in their native tissue environment and highlight the potential of spatial lipidomics to reveal metabolic dependencies in host-microbe associations.

## Introduction

Members of the family *Endozoicomonaceae* establish close associations with a wide range of marine animals including corals, sponges, fish, and bivalves. Their interactions span a continuum from mutualistic to parasitic lifestyles [1]. Species of the genus *Endozoicomonas* are among the most prevalent bacterial symbionts in corals and sponges and are thought to benefit the host through involvement in nitrogen and phosphorus cycling as well as provision of essential amino acids and B vitamins [1, 2]. In addition, a small number of parasitic associations has been described.

*Candidatus* Endonucleibacter childressi is an example for a parasitic bacterium belonging to the family *Endozoicomonadaceae*. It infects the nuclei of epithelial cells in the gill tissue of the deep-sea mussel *Gigantidas childressi* and replicates to as many as 80,000 bacterial cells within a single host nucleus. A dual transcriptomic study suggested that *Ca*. Endonucleibacter heavily exploits lipids as a carbon and energy source [3]. The parasite actively secretes a lipase to degrade host lipids and imports fatty acids via transporters. In response, the host cell increases the expression of enzymes and transporters involved in lipid, amino acid and oligosaccharide metabolism to compensate for the depletion of its intracellular storage compounds [3]. It remains unknown to what extent infection induced an altered metabolic phenotype of the host cell or if the exploitation of host lipids is reflected in the parasite’s metabolic phenotype.

Spatial metabolomics using mass spectrometry imaging technique is uniquely suited to capture such metabolic changes *in situ* [4]. Matrix-assisted laser desorption/ionization mass spectrometry imaging (MALDI-MSI) is often used to study host-pathogen interactions from a metabolic perspective. It enables the visualization of hundreds of metabolites at micrometer resolution directly within the infected tissue [5–9]. MSI spatial resolution (5-20 µm) remains insufficient to profile single bacterial cells yet a detection of bacterial membrane lipids in colonized tissue regions can identify metabolic markers for symbiont-associated or infected host tissues [10–12].

By combining MALDI-MSI with 16S rRNA fluorescence *in situ* hybridization (16 S rRNA FISH), a method to visualize bacteria in tissue sections using DNA-based probes, changes in the spatial metabolome can be linked to the presence of bacteria. The combination of MALDI-MSI and 16S rRNA FISH has been applied previously to symbiotic systems to study microbial metabolism *in situ* [12–14]. It was shown that this approach is capable of identifying lipid biomarkers for intracellular, non-culturable symbionts as well as linking changes in the host metabolome to the presence of bacterial symbionts [12]. For the methane-oxidizing (MOX) symbionts that co-occur with *Ca*. Endonucleibacter childressi in the gill tissue of *G. childressi*, hopanoids have previously been identified as bacterial lipid biomarkers. In addition, symbiont-hosting tissue regions have been associated with high triglyceride abundance [12].

Here, we combine MALDI-MSI with 16S rRNA FISH to map lipidomic changes in epithelial cells infected with *Ca*. Endonucleibacter childressi. We investigate whether infection leads to an altered lipidomic phenotype of the host cell and focus on the phospholipid profile of *Ca*.

Endonucleibacter childressi in its native tissue environment. The co-occurrence of *Ca. Endonucleibacter* and the mussel’s chemosynthetic symbionts in spatially distinct niches within the gill provides a unique opportunity to compare the lipidomes of parasitic and mutualistic intracellular gammaproteobacteria within the same host.

## Materials & Methods

### Sample collection

Samples were taken during the cruise E/V Nautilus in 2015 at sampling side MC853 (28°7’24.96”N, 89°8’23.639”W, depth 1071m). The mussels were retrieved from the vent side using a remotely operated vehicle and transported to the sea surface in an insulated container. The gill tissue was dissected immediately after retrieval of the mussels, snap frozen in liquid nitrogen and stored at -80°C. The gills of different *Gigantidas childressi* specimens were used for MALDI-MSI and LC-MS/MS analyses.

### MALDI-MSI

The frozen gill tissue of one specimen was embedded in 2% carboxymethylcellulose gel (molecular weight ∼700.000 g*mol^-1^), snap-frozen in liquid nitrogen and then sectioned into 12 μm thick tissue sections using a cryotome (Leica CM3050 S, Leica Biosystems) with a chamber temperature of -21°C and object holder temperature of -19°C. Sections were thaw-mounted onto polysine slides (Thermo Scientific). Overview images of the tissue sections were acquired with an Olympus BX53 microscope. Regions of interest were defined and marked with white paint as described previously [10]. A solution of super 2,5-dihydroxybenzoic acid (Sigma-Aldrich, 99% purity, 30 mg*ml^-1^ in acetone:water (6:4 (v:v) with 0.1% trifluoroacetic acid)) was prepared and 225 μL of SDHB solution were deposited onto each slide using a pneumatic sprayer system (SMALDIPrep, TransMIT) using the following parameters (flow rate = 7.5 μl*min^-1^, rotation = 350 rpm, N_2_ flow = 5 l*min^-1^).

MSI datasets were acquired on a MALDI-MSI system consisting of an atmospheric pressure ion source (AP-SMALDI 10, TransMIT) coupled to a Q Exactive HF mass spectrometer (ThermoFisher Scientific). The instrument was calibrated on SDHB ions and laser focus was optimized using a red marker pigment before each measurement [10]. The two datasets MPIMM_192 and MPIMM_193 were acquired from gill tissue sections of the same gill tissue specimen. Measurements were conducted in positive ion mode covering a mass range of 300-1200 Da and 150-1200 Da respectively at a mass resolving power of 240.000 at *m/z* 200.

### Fluorescence microscopy using 16S rRNA FISH probes

16S rRNA FISH probes were used to specifically label bacteria at the species level as previously described [10]. After MALDI-MSI, the matrix was removed and the tissue was fixed for microscopy by submerging the slide in a 2% paraformaldehyde/phosphate-buffered saline (PBS) solution for one hour followed by two washing steps with PBS and a final washing step in ethanol 96% (v:v). A liquid blocker (PAP-Pen, Science Services) was applied to encircle the region of interest and hold the hybridization mixture containing the 16S rRNA FISH probes in place. A Cy3-labelled 16S rRNA probe specific for *Ca*. Endonucleibacter (5’GCTAGACCTGTTACCGCT’3) and a atto647-labelled EUB-338 probe (5’GCTGCCTCCCGTAGGAGT’3) were used to visualize the localization of bacteria in the gill tissue [15, 16]. Hybridization of tissue sections was conducted in a formamide- and water-saturated atmosphere at 46°C for 2 hours. Afterwards, tissue sections were washed with washing buffer, dipped in PBS and then ethanol 96% (v:v). A 4′,6-diamidino-2-phenylindole (DAPI) stain (10 min) was conducted to visualize host and bacterial DNA. A negative control was used to check for unspecific binding on a consecutive tissue section. Tissue sections were mounted with VECTASHIELD®. Fluorescence microscopy images were acquired on an Olympus BX3. Images were processed in ImageJ by adjusting brightness and contrast.

### Statistical analyses of MALDI-FISH datasets

MSI datasets and corresponding FISH images were imported to SCiLS lab (Bruker Daltonics, v.2024a Pro) and overlayed using *m/z* 682.4773 and *m/z* 848.5052 as ions to outline the filaments and infected cells. Regions of interest in the MSI datasets were defined manually based on the fluorescence signals (DAPI for the host tissue, *Ca*. Endonucleibacter-specific probe for infected regions and EUB probe for symbiont regions).

Features highly abundant in infected regions were identified manually and using a colocalization analysis (*m/z* 754.4783 was used as a reference ion, highly abundant in infected regions). After a manual quality check 41 features were identified, exported and then grouped using the mass2adduct package in R to identify different adducts of the same metabolite [17]. Features were categorized grouping together all adducts of metabolites based on accurate mass differences (**Table S1**).

Intensity values for potassium adducts of the five lipid biomarkers in infected, uninfected and symbiont regions were exported from SCiLS. Pseudocounts were created using the half minimum nonzero value and log2-normalized to plot intensity values. Hierarchical clustering using Euclidean distances and plotting was done with the pheatmap package in R [18].

Relative abundance of triglycerides was compared between 10 representative infected and 10 uninfected regions for all features putatively annotated as triglycerides using the METASPACE platform. Intensity values for the features were extracted using SCiLS and significance was calculated using a two-sided, two sample t-test in R with Benjamini-Hochberg correction.

Fluorescence intensity for the *Ca*. Endonucleibacter-specific 16S rRNA FISH probe in infected and uninfected regions was extracted using ImageJ. TIC-normalized signal for the five biomarkers was extracted for the same regions from the MSI data using SCiLS. Mean fluorescence per region and average signal intensity of each biomarker per region was then used to create a scatterplot. Pearson correlation coefficients were calculated in R to determine correlation of fluorescence intensity of the 16S rRNA probe specific to *Ca*. Endonucleibacter and MALDI-MSI signal for the different lipid biomarkers.

### Bulk metabolomics using LC-MS/MS

#### Sample preparation

Lipids were extracted from small pieces (50-100 mg) of gill tissue of four *Gigantidas childressi* specimens using a biphasic methanol-chloroform extraction protocol. The tissue was placed in screw cab tubes with silica beads (1.1-1.2 mm diameter, Sigmund Linder) and 8 µL ice-cooled methanol was added per milligram of wet tissue. Tissues were homogenized using a FastPrep-24 5G (MP Biomedicals, 2x10 seconds burst at 6.5 m*s^-1^) and transferred into a 3 mL exetainer containing 8 µL of ice-cooled chloroform per milligram tissue. After vortexing the exetainers for 15 seconds, they were placed on ice and 7.2 µL mg^-1^ tissue HPLC grade water was added to each exetainer. The exetainers were vortexed again for 30 seconds, put on ice for 10 minutes to achieve phase separation. Tissue debris was removed by centrifugation (10 minutes, 4°C, 2500x g). The tubes were left at room temperature for 10 minutes to achieve phase separation. The chloroform phase was transferred to a HPLC-MS vial (1.5-HRSV 9mm Screw Thread Vials, Thermo Fisher™) and evaporated to dryness under nitrogen. The dried extracts were stored at -80°C until further analyses. Samples were reconstituted in 500 μl chloroform before analysis. The reconstituted extract was diluted 1:10 in acetonitrile:methanol:water (1:1:1 (v:v:v)) and transferred to an HPLC-MS vial. Aliquots of diluted samples were pooled as quality control samples. All organic solvents were LC-MS grade: acetonitrile (ACN; Honeywell, Honeywell Specialty Chemicals), isopropanol (IPA; BioSolve), formic acid (FA; Sigma-Aldrich) and methanol (MeOH, BioSolve). Water was deionized using the Astacus MembraPure system (MembraPure).

#### High-resolution LC-MS/MS

Samples were measured on a Q Exactive Plus Orbitrap (Thermo Fisher Scientific) equipped with a HESI probe and a Vanquish Horizon UHPLC system (Thermo Fisher Scientific). Lipids were separated on an Accucore C30 column (150 x 2.1 mm, 2.6 μm, Thermo Fisher Scientific) at 40°C using a solvent gradient (**Table S2**) with buffer A (60:40 acetonitrile:water (v:v), 10 mM ammonium formate, 0.1% formic acid) and buffer B (90:10 isopronaol:acetonitrile (v:v) 10 mM ammonium formate, 0.1% formic acid). A flow rate of 350 μl min^-1^ was used.

The injection volume of the sample was 10 μl. MS measurements were acquired in positive and negative ion mode in two separate runs for a mass detection range of *m/z* 150-1500. Mass resolution was set to 70.000 for MS scans and to 35.000 for MS^2^ scans at *m/z* 200. MS/MS scans of the eight most abundant precursor ions were acquired in both ionization modes excluding the 200 most abundant *m/z* detected in the extraction blank. Dynamic exclusion was enabled for 30 s and collision energy was set to 30 eV (more details on the MS method in **Table S3**).

#### Data analysis

LC-MS/MS files were centroided and converted to mzML format using MSConvert (ProteoWizard) [19]. Raw data were inspected using Freestyle (v.1.6.75.20). Extracted ion chromatograms (EICs) for adducts of the five biomarkers (**Figures S1-S5)** were extracted and fatty acid composition was determined manually by extracting MS/MS spectra for [M-H]^-^ adducts. EICs and MS/MS spectra were extracted and plotted with MSnbase (v2.30.1) in R [20].

Data preprocessing for molecular network analysis was performed in R using the XCMS (v4.0.2), MSnbase (v2.30.1), and CAMERA (v1.56.0) packages [20–23]. Positive and negative ion mode datasets were processed separately. Briefly, peak detection was performed using the CentWave algorithm, retention time alignment was conducted using Obiwarp and chromatographic peak grouping was performed based on peak density (**Table S4**). Missing features were imputed and isotopes and adducts were annotated. MS/MS spectra associated with detected features were extracted, normalized to base peak intensity and filtered to retain only spectra containing characteristic phospholipid head group fragments (ethanolamine-phosphate (*m/z* 140.0118, negative mode) and phosphocholine (*m/z* 184.0733, positive mode) using a mass tolerance of 0.005 Da). Spectra where the base peak did not match the precursor ion were discarded. Fragment ions with relative intensities below 5% of the base peak were removed to reduce spectral noise. The resulting filtered MS/MS data were used to generate a list of putative phosphatidylcholines (PCs) and phosphatidylethanolamines (PEs). These MS/MS-derived molecular ions were then matched to MALDI features within a ± 5 ppm mass window. Because LC-MS/MS data were acquired in both ionization modes but MALDI-MSI data were collected in positive mode only, an additional step was required for features detected in negative ion mode. For these, putative positive-mode adducts ([M+H]^+^, [M+Na]^+^, [M+K]^+^) were calculated and the resulting theoretical *m/z* values were then matched to MALDI features. Matched ions were used to refine annotations for nodes of PCs and PEs in molecular networks.

### Molecular network analysis

From the MSI data, tissue-related features were extracted with the CatBoost package in R [24]: A model was trained and validated using a representative subset of 5% of all features respectively. Features of the training and validation dataset were manually classified into tissue-related features and background based on the spatial distribution. All features showing signal co-localizing with host tissue and little to no signal outside the tissue section were classified as tissue-related. Remaining features were classified as background signal. Next, TIC-normalized intensity was extracted for all features in nine representative regions of the MALDI image covering different tissue and background areas. Based on signal intensity of the features in the different regions Catboost was trained and then used to classify the remaining features. Out of 16456 features 1235 were classified as tissue-related. For a representative subset of 5% of the classified features the accuracy was above 95% for the trained model (sensitivity: 68.1%, specificity: 99.6%). The identified tissue-related features were used to create molecular networks.

MS^1^ networks were created in Cytoscape using the app MetaNetter2. A peak list comprising all tissue-related features after removal of 13C isotopes (n=1000) was imported and used to create nodes. A list of common mass transformations was used to connect the nodes (**Table S5**). Nodes were color-coded according to the lipid class annotations obtained from the METASPACE platform [25]. Whenever possible, annotations for PCs and PEs were refined based on the LC-MS/MS data as described above. For the PE network shown in Figure 3B, annotations were revised manually based on fragmentation patterns in LC-MS/MS data.

### Transcriptomics

#### Expression analysis of ‘*Ca*. E. childressi’

Transcriptomics analyses were done as in Porras et. al 2024 [3]. In short, RNA reads (SRX24174227) were quality-trimmed and adaptors removed using BBDuk v38.90 (https://sourceforge.net/projects/BBMap). To remove non-mRNA contaminants, reads were mapped against the rRNA and tRNA SILVA database v132 using BBMap v38.90 (identity: 0.85) [26]. The expression of *Ca*. Endonucleibacter childressi was quantified using Kallisto v.0.44.0 with default parameters using *Ca*. Endonucleibacter childressi (GCA_030674875.1) as reference genome [27]. Transcription levels were normalized to the single-copy housekeeping gene *RecA* and mapped onto metabolic pathways using Pathway tools v13.0 for the reconstruction of *Ca*. Endonucleibacter childressi lipid metabolism [28].

### Functional annotation

#### Protein domain analysis

To verify the genes considered for the reconstruction of *Ca*. Endonucleibacter childressi lipid metabolism, Hidden Markov Model (HMM) and protein domain analyses were applied. HMM profiles of publicly available *Endozoicomonadaceae* protein sequences analogous to those to be verified in *Ca*. Endonucleibacter childressi were generated. Then, hmmsearch (http://hmmer.org/) was performed against the profiles using default thresholds (E-value 1x10E-3) to identify and functionally annotate the investigated genes in *Ca*. Endonucleibacter childressi. The protein domains of hmmsearch hits were double checked with the NCBI protein domain search platform (https://www.ncbi.nlm.nih.gov/Structure/cdd/wrpsb.cgi). Only those hmmsearch hits which protein domains congruent with their functional annotation were considered for the reconstruction of lipid metabolism.

## Results & Discussion

### The lipid profile of *Ca*. Endonucleibacter childressi infected cells is dominated by bacterial glycerophospholipids

We conducted MALDI-MSI and 16S rRNA FISH on gill tissue sections obtained from a *Gigantidas childressi* specimen from the Gulf of Mexico. By combining these two techniques, we visualized the spatial lipidome and microbiome on the same tissue sections to identify lipidomic features associated with the infection of *Ca*. Endonucleibacter childressi (**Figure 1**). Spatial lipidomics was performed at 10 μm pixel size and revealed hundreds of tissue-related mass spectrometry features. The majority of features was putatively annotated as glycerophospholipids or glycerolipids using the MSI annotation platform METASPACE [25]. The spatial microbiome was visualized using a 16S rRNA probe specific to *Ca*. Endonucleibacter childressi and a probe labelling all bacteria (eubacterial probe, EUB). Cells infected with *Ca*. Endonucleibacter childressi were confined to epithelial cells in the outer edge of the tissue and spatially distinct from the symbiont-colonized epithelial cells along the gill filaments [3]. Uninfected regions were present in the same tissue section and served as internal reference regions for uninfected epithelial cells in our analyses (**Figure 1**).

**Figure 1:**
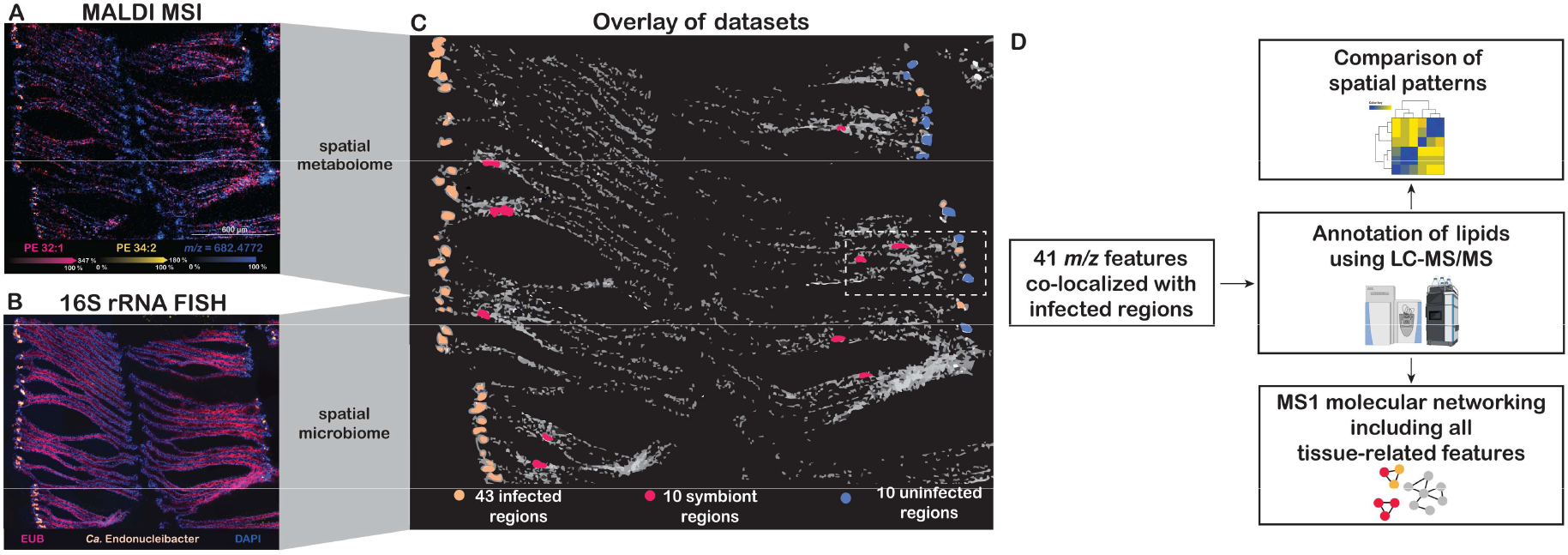
Identification of lipid biomarkers for *Ca*. Endonucleibacter childressi *in situ* by combining spatial lipidomics and microscopy with fluorescence *in situ* hybridization using 16S rRNA FISH probes. **A)** Spatial distribution of exemplary lipids in the gill tissue. The lipid PE 32:1 (magenta, [M+K]^+^, *m/z* 728.4629 ± 3 ppm) is found across symbiont and infected regions, PE 34:2 (yellow, [M+K]^+^, *m/z* 754.4789 ± 3 ppm) is specific to *Ca*. Endonucleibacter childressi infected regions and a third, unannotated feature (blue, *m/z* 682.4772 ± 3 ppm) outlines the host tissue. **B)** Localization of bacterial cells in gill tissue sections using 16S rRNA FISH probes. A eubacterial probe (magenta) was used to label all bacteria. A specific 16S rRNA FISH probe (yellow) shows the localization of the parasitic bacterium *Ca*. Endonucleibacter childressi. The DNA stain DAPI was used to visualize the outlines of the host tissue (blue). **C)** MALDI-MSI and FISH datasets were overlayed to compare the spatial metabolome of different regions of the gill tissue. *Ca*. Endonucleibacter childressi infected regions (43 ROIs) were defined based on the FISH signals. Infected cells in the outer edge of the tissue occurred in small groups of two to five individual host cells and were grouped into regions of interest (ROIs). Grouping helped to circumvent resolution constraints and limit uncertainties during correlation of MSI and FISH images. Representative symbiont and uninfected reference tissue regions (each 10 ROIs) were defined for further analyses. Dotted lines indicate area displayed in Figure 2. **D)** Lipid analysis workflow for the identification of features characteristic for infected regions. Features were annotated using LC-MS/MS data from bulk tissue extracts. Spatial patterns of lipid biomarkers were compared and MS1 molecular networks were created.

To identify lipidomic changes in *Ca*. Endonucleibacter childressi infected host cells, we used co-localization analyses after co-registration of the MSI and FISH dataset (**Figure 1A-C**). These revealed 41 *m/z* features highly abundant in infected regions (**Table S1**). Features were grouped using the mass2adduct package and annotated using the METASPACE platform [25].

MALDI-MSI revealed a strong enrichment of four glycerophosphoethanolamines (PE) and one glycerophosphoglycerol (PG) in *Ca*. Endonucleibacter childressi infected regions **(Table 1, Figure 2)**. Based on accurate mass, these features were initially annotated as PE 32:1, PE 34:4, PE 34:2, PE 36:5, and PG 36:5 (**Table 1**). To validate and refine these annotations, we performed untargeted LC-MS/MS lipidomics on gill tissue extracts from four *G. childressi* specimens. LC-MS/MS confirmed the presence of isomers for the PE species and resolved their fatty acid composition. The PEs consisted of even-chain C16 and C18 fatty acids with varying degrees of unsaturation ranging from no to four double bonds per fatty acid (**Table 1, Figures S2-S5**).

**Table 1:** Five glycerophospholipids were detected in *Ca*. Endonucleibacter childressi infected gill tissue regions. Potassium adducts for all lipids are used as representative features for the five biomarkers in the MSI datasets throughout the manuscript. Annotations obtained via METASPACE [25] were revised using LC-MS/MS data of bulk tissue extracts. For example, features annotated as PE 34:4 or PC 31:4 were found to represent at least three isomeric PEs: PE 18:1_16:3, PE 18:3_16:1, and PE 18:4_16:0 were detected indicating the presence of multiple lipid species sharing the sum formula PE 34:4 in the gill tissue. Similarly, LC-MS/MS analyses confirmed that the remaining three putative PC/PE features were all PEs, each composed of even-chained C16 and C18 fatty acids with varying degrees of unsaturation (**Figures S2-5**). Features corresponding to the putatively annotated PG 36:5 were detected with LC-MS yet their abundance was too low to obtain MS/MS spectra and refine the annotation (**Figures S1**).

| MALDI-MSI |  |  |  |  | LC-MS/MS |
| --- | --- | --- | --- | --- | --- |
| Sum composition (LipidMaps) | adduct | *m/z* observed | *m/z* theoretical | Δ ppm | confirmed lipid species |
| PE 32:1 | [C<sub>37</sub>H<sub>72</sub>NO<sub>8</sub>P+K]<sup>+</sup> | 728.4629 | 728.4627 | 0.27 | PE 16:0_16:1 |
| PE 34:4 | [C<sub>39</sub>H<sub>70</sub>NO<sub>8</sub>P+K]<sup>+</sup> | 750.4458 | 750.4471 | 1.73 | PE 18:1_16:3,<br>PE 18:3_16:1,<br>PE 18:4_16:0 |
| PE 34:2 | [C<sub>39</sub>H<sub>74</sub>NO<sub>8</sub>P+K]<sup>+</sup> | 754.4788 | 754.4784 | 0.53 | PE 18:1_16:1,<br>PE 18:2_16:0 |
| PE 36:5 | [C<sub>41</sub>H<sub>72</sub>NO<sub>8</sub>P+K]<sup>+</sup> | 776.4618 | 776.4627 | 1.16 | PE 18:4_18:1,<br>PE 18:3_18:2 |
| PG 36:5 | [C<sub>42</sub>H<sub>73</sub>NO<sub>10</sub>P+K]<sup>+</sup> | 807.4560 | 807.4573 | 1.61 | - |

**Figure 2:**
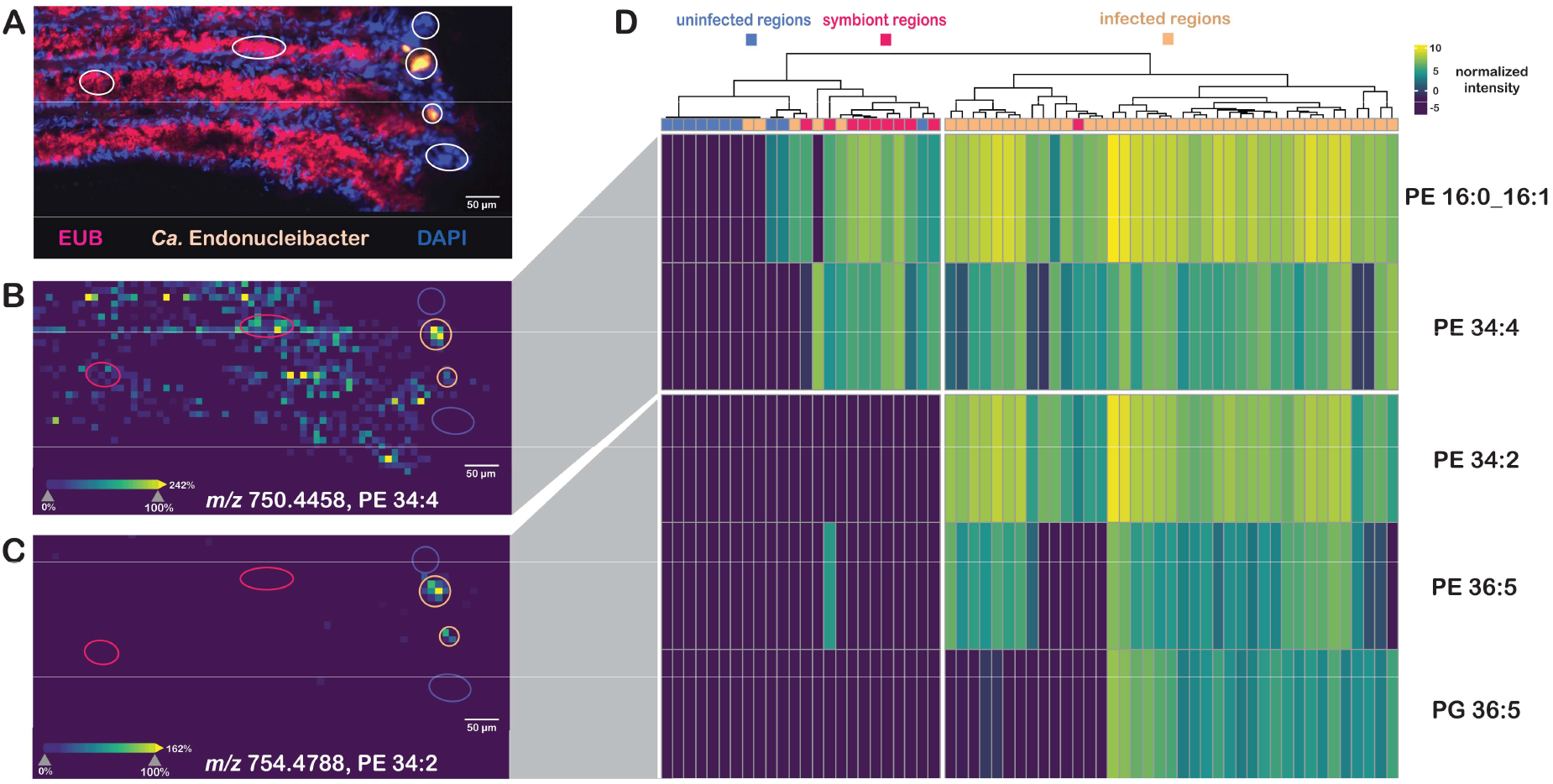
Lipid biomarkers show two spatial patterns in *Ca*. Endonucleibacter childressi infected gill tissue. **A)** Zoom-in of the gill tissue showing the localization of all bacteria (magenta, EUB probe), *Ca*. Endonucleibacter childressi (yellow, *Ca*. Endonucleibacter probe) and host tissue (blue, DAPI stain). MALDI-MSI images of the same tissue area show the two spatial patterns that were observed: **B)** Lipid distribution matching presence of all bacteria, e.g. PE 34:4 ([M+K]^+^, *m/z* 750.4458 ± 3 ppm). LC-MS/MS analyses of tissue extracts showed that PE 34:4 comprises at least three isomeric species (**Table 1**). Because isomers cannot be distinguished by our MSI approach, the observed PE 34:4 signal represents the combined spatial distribution of all isomers. It remains unclear whether all PE 34:4 isomers are shared between *Ca. Endonucleibacter childressi* and the MOX symbiont or whether some are specific to one of the two species. **C)** Lipid distribution co-localizing with *Ca*. Endonucleibacter childressi infected regions, e.g. PE 34:2 ([M+K]^+^, *m/z* 754.4788 ± 3 ppm). Regions included in analysis are indicated in orange (*Ca*. Endonucleibacter childressi infected), magenta (symbiont regions) and blue (uninfected host region). **D)** The heatmap displays lipid biomarker signal across regions in the analyzed gill tissue section as log2normalized intensity.

### Glycerophospholipid profiles of *Ca*. Endonucleibacter childressi and MOX symbionts partially overlap

Examination of the spatial distribution of the identified lipid biomarkers revealed two localization patterns: Three lipids, PE 34:2, PE 36:5 and PG 36:5, were exclusively detected in *Ca*. Endonucleibacter childressi infected regions making them specific biomarkers for infected regions of the gill tissue (**Figure 2A-B**). In contrast, PE 16:0_16:1 and PE 34:4 were detected not only in infected regions but also along the gill filaments colonized by the mussel’s mutualistic symbionts (**Figure 2A+C, Figures S6-7, Table S6-S7**).

To compare the relative abundance of the five bacterial biomarkers in infected, uninfected and symbiont regions, we used hierarchical clustering of all the ROIs (**Figure 2D**). The majority of *Ca*. Endonucleibacter childressi infected regions clustered together due to high abundance of the five bacterial glycerophospholipids. Uninfected reference regions and symbiont ROIs formed each a separate cluster. The symbiont cluster is characterized by low abundance of the gammaproteobacterial biomarkers PE 16:0_16:1 and PE 34:4 and absence of the three *Ca*. Endonucleibacter childressi specific lipids.

A few *Ca*. Endonucleibacter childressi infected ROIs with positive FISH signal displayed lipid profiles similar to those of both symbiont and uninfected regions. This discrepancy likely reflects differences in sensitivity of the two methods to detect bacteria, with FISH being more sensitive to low bacterial density than MALDI-MSI. The variable signal of lipid biomarkers across infected regions may reflect differences in bacterial abundance and, potentially, different stages of infection. Consistent with this interpretation, lipid signal intensity was positively correlated with *Ca*. Endonucleibacter childressi 16S rRNA FISH signal intensity in infected regions, although quantitative interpretation is currently limited by the lack of standardized approaches for estimating bacterial biomass from either FISH or MALDI-MSI data (**Supplementary Text, Figure S8, Table S8**).

We hypothesize that the five glycerophospholipids are bacterial membrane lipids. We observed a high degree of correlation between the spatial distribution of the five lipids and tissue regions colonized with bacteria. PEs and PGs dominate the polar lipid profile of close relatives of the MOX symbionts and *Ca*. Endonucleibacter childressi and are likely the most abundant classes of glycerophospholipids in the two host-associated gammaproteobacteria [1, 29, 30]. Based on our MSI data, we cannot conclusively assign these lipids to the bacteria because the signals originate from infected tissue containing both host and bacterial biomass. However, their spatial association with bacterially colonized regions and the expected phospholipid composition as seen in close relatives, supports their interpretation as bacterial phospholipids. Overall, our findings provide a first characterization of the dominant glycerophospholipids of *Ca*. Endonucleibacter childressi and the MOX symbionts in their native tissue environment and reveal a partially shared phospholipid profile between these two host-associated gammaproteobacteria.

### Host glycerolipid content remained unaffected by infection

Previous investigations based on transcriptomic data inferred that *Ca*. Endonucleibacter childressi uses host lipids as a nutritional resource for its own survival and replication. We explicitly compared the signal intensity of glycerolipids in infected and uninfected cells but found no difference in relative abundance in infected and uninfected regions. Triglycerides showed highest signal throughout symbiont-colonized filaments (**Figure S9**). Little signal was observed in the outer edge of the gill tissue where infection occurs and no significant change in glycerolipid signal could be detected with the semi-quantitative MSI approach. Possible explanations for these observations can be that infection increases host lipid turnover without substantially altering the overall glycerolipid content of the infected cells indicating that the host cell is capable of compensating for the use of storage lipids. Alternatively, *Ca*. Endonucleibacter childressi may utilize other host lipid resources, such as free fatty acids or phospholipids. Further studies combining laser-capture microdissection with quantitative LC-MS/MS lipidomics may provide a more sensitive assessment of lipid dynamics during *Ca*. Endonucleibacter childressi infection [32].

### Bacterial and host PEs cluster together in molecular networks

To study the chemical relationship of the five identified bacterial biomarkers with the overall lipidome of the mussel’s gill tissue, we created molecular networks. These networks show individual ions (MS1) as nodes that are connected through hypothetical but common molecular transformations allowing us to explore chemical relationships between known and unknown metabolites within a dataset. Tissue-related ions were extracted from the MSI dataset and used as network nodes. Five common lipid modifications were used to cluster nodes into subnetworks.

The resulting network is dominated by glycerophospholipids with glycerophosphocholines (PC) and PEs forming the largest subnetworks (**Figure 3A**). This dominance is expected as PCs and PEs are the major membrane lipids reported for close relatives of the mussel host and the gammaproteobacteria [1, 33]. PCs originate exclusively from the host whereas PEs can be synthesized by the host and the two gammaproteobacteria. The networks allowed us to identify further putative PGs, phosphatidic acids (PAs) and triglycerides whereas most nodes remained unannotated. These ions likely represent metabolites that are not covered by the LipidMaps database used for annotation as well as matrix adducts or in-source fragments.

**Figure 3:**
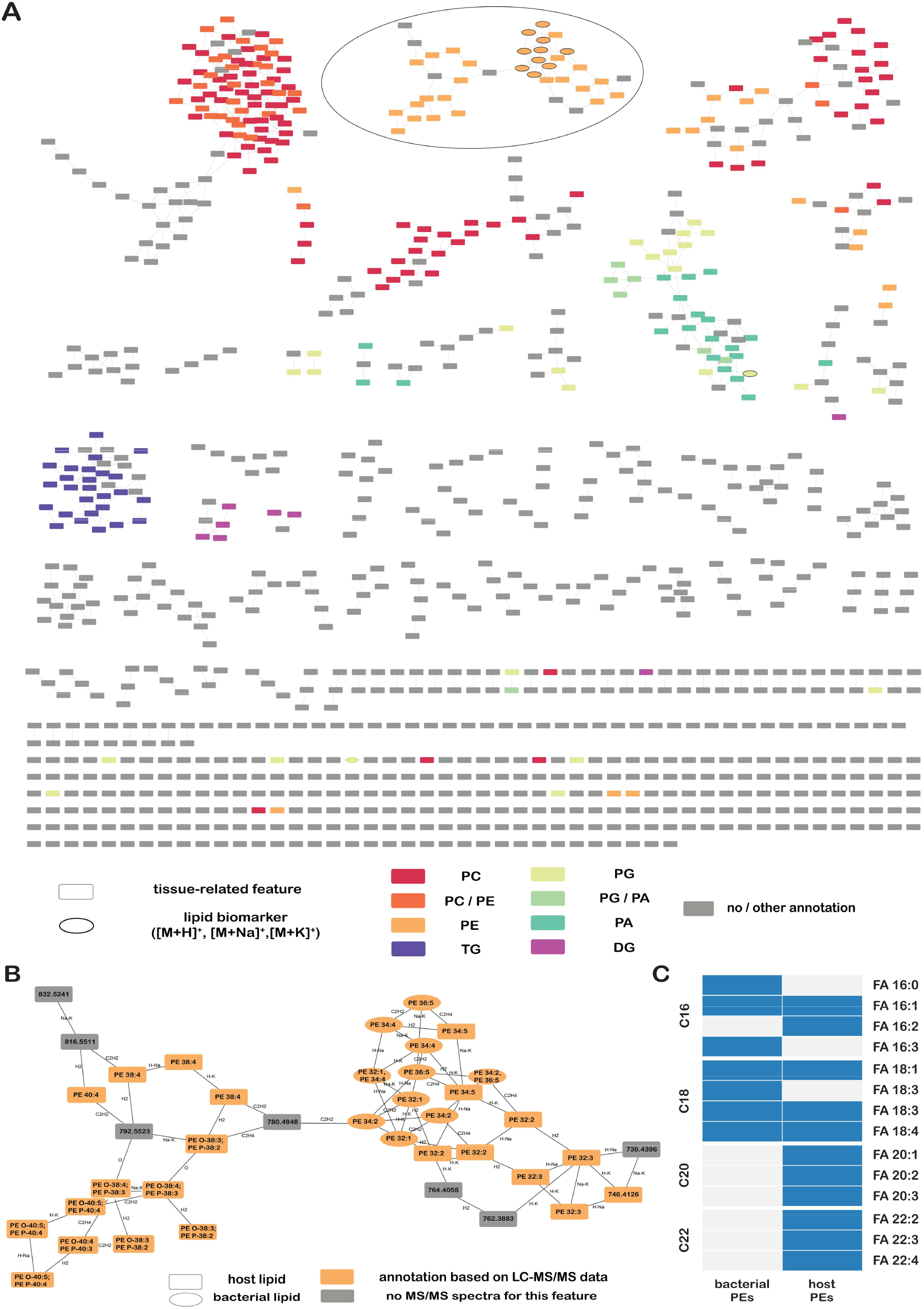
Molecular networking identifies similar host and bacterial PEs which were used for subsequent fatty acid composition analysis. A) Molecular networks visualizing the chemical diversity of the spatial lipidome of the gill tissue. Colored nodes represent annotated features. Edges show common mass differences for lipids. Proton and alkali adducts of biomarkers characteristic for *Ca*. Endonucleibacter childressi infected regions are shown as ellipses. The largest PE subnetwork containing both bacterial and host PEs shown in Figure 3B is highlighted. B) Subnetwork containing bacterial and various host PEs. PEs were assigned either bacterial or host origin depending on their spatial distribution in the gill tissue. Features co-localizing with the DAPI stain and little to no co-localization with the eubacterial probe were classified as host lipids. PEs previously identified as highly abundant in bacterial regions were highlighted as ellipses. Annotations were revised based on LC-MS/MS spectra (**Table S9**). C) The fatty acid profile of bacterial and host PEs partially overlaps. Fatty acids were identified based on accurate mass in LC-MS/MS spectra of features identified as bacterial and host PEs. Blue indicates presence and white absence.

In our following analyses, we focused on the largest PE subnetwork containing the four bacterial PEs as well as various host PEs (**Figure 3B**). All PE species within the network had an even total number of carbon atoms across their two fatty acyl chains. To find out more about the fatty acid composition and potential differences between host and bacterial PEs, we used our LC-MS/MS data to manually curate annotations for all features within the PE network (**Figure 3B**).

### Polyunsaturated fatty acids suggest host influence on bacterial lipid composition

The PEs assigned to *Ca. Endonucleibacter* childressi and the MOX symbionts exhibited highly similar fatty acyl compositions. Bacterial PEs consisted of combinations of C16 and C18 fatty acids with no to up to four double bonds (**Figure 3C, Table S9**), suggesting that the gammaproteobacteria share a largely similar fatty acid pool. The fatty acyl composition of host PEs partially overlapped with the bacterial lipids, as they likewise contained numerous C16 and C18 fatty acids. However, the host fatty acid pool was more diverse and additionally included C20 and C22 fatty acids which were not detected in bacterial PEs (**Figure 3C**). Overall, the fatty acid composition was consistent with previous lipidomics studies on gill tissue extracts of *Bathymodiolus* mussels [34], which reported predominantly even-chain fatty acids (C16-C22) with varying degrees of unsaturation from polar lipid-derived fatty acid analyses.

Polyunsaturated fatty acids such as FA 16:3, FA 18:3 and FA 18:4 have not been reported previously from cultured, close relatives of *Ca*. Endonucleibacter childressi and the MOX symbionts. For *Endozoicomonas* and *Parendozoicomonas* strains, close relatives of *Ca*. Endonucleibacter childressi, C16 and C18 fatty acids containing no or a single double bond were reported to be the most abundant fatty acids making up the polar lipid fraction [1]. Similarly, the fatty acid profile of the free-living methanotroph *Methyloprofundus sedimentii*, a close relative of the mussel’s MOX symbionts, was reported to have a fatty acid profile dominated by saturated, mono- and di-unsaturated C16 and C18 fatty acids [30]. To our knowledge, fatty acids with more than two double bonds have not been reported from cultured, close relatives [1, 30].

*Ca*. Endonucleibacter childressi and the MOX symbionts themselves are currently unculturable so it remains unknown whether the occurrence of polyunsaturated fatty acids is a lineage-specific feature of these gammaproteobacteria or if this lipidomic feature is associated with growth in the host environment. Our hypothesis is that these fatty acids are derived from metabolic interactions with the host, potentially through the uptake and subsequent remodeling of polyunsaturated, host-derived fatty acids. Further studies including lipid transfer experiments would be needed to prove this hypothesis and determine the metabolic origin of the polyunsaturated fatty acids detected in the putative bacterial phospholipids.

### *Ca*. Endonucleibacter childressi shows capabilities for uptake and remodelling of host-derived fatty acids

To investigate the potential for utilization and modification of host-derived lipids, we reanalyzed the available transcriptomic dataset for *Ca*. Endonucleibacter childressi with a focus on lipid metabolism. We found that it encodes and expresses a complete set of genes involved in fatty acid uptake, β-oxidation, fatty acid biosynthesis and remodeling (**Figure S11**). The high expression of β-oxidation genes (e.g. *fadA, fadB, fadE*) suggests that host-derived fatty acids are an important source of carbon and energy. At the same time, expression of fatty acid biosynthesis and remodeling genes (e.g. *fabD, fabZ, fabI, fabB, fabG*) indicates that imported fatty acids can be modified before incorporation into membrane lipids. Although genes involved in phospholipid synthesis (*plsB, pssA, psd*) were expressed at comparatively low levels and transcripts for *plsC* and *pgpA* were not detected in the available dataset, the combined genomic and transcriptomic evidence suggests that *Ca*. Endonucleibacter childressi is capable of incorporating and remodeling host-derived fatty acids for phospholipid synthesis.

Comparable lipid acquisition and utilization strategies have also been proposed from genomic analyses of the mutualistic coral symbiont *Endozoicomonas* associated with *Acropora loripes* [35], suggesting that host-derived fatty acids may commonly contribute to the fatty acid pool of these host-associated bacteria. Whether the same mechanisms underlie the occurrence of polyunsaturated fatty acids in the phospholipid PE 34:4 that was also assigned to the MOX symbionts remains unknown, but the observed fatty acid profile observed here raises this possibility.

## Conclusion

Here, we determined the phospholipid profile of the intranuclear parasitic bacterium *Ca*. Endonucleibacter childressi directly within its native host tissue using spatial lipidomics and 16S rRNA FISH. We observed a partial overlap in the phospholipid profile with the mussel’s chemoautotrophic symbionts showing that there are shared lipidomic features between the two gammaproteobacteria despite different lifestyles and modes of carbon and energy acquisition.

Phospholipids assigned to *Ca. Endonucleibacter childressi* contained polyunsaturated fatty acids, including FA 16:3, FA 18:3 and FA 18:4, which have not been reported from cultured relatives [1, 30]. Whether these polyunsaturated fatty acids represent lineage-specific features or reflect a host-associated lifestyle characterized by acquisition and remodeling of host-derived fatty acids remains unresolved. In *Ca. Endonucleibacter childressi*, the expression of genes involved in fatty acid uptake and remodeling suggests that host-derived lipids contribute to its fatty acid pool. Similar lipid scavenging strategies have been proposed for mutualistic *Endozoicomonas* [30], suggesting that the acquisition and remodeling of host-derived fatty acids may represent a common metabolic strategy among host-associated members of the *Endozoicomonodaceae*.

This study highlights the value of characterizing host-microbe associations in their native tissue environment using spatial analyses. Our findings suggest that lipid profiles of host-associated bacteria can provide insights into metabolic interactions between hosts and their symbionts. Comparative studies across mutualistic and parasitic systems will be essential to determine the extent to which genomic potential, lifestyle, and host context shape bacterial lipid profiles.

## Acknowledgments

We thank the Symbiosis Department of the Max Planck Institute for Marine Microbiology for scientific discussions, the crew and captains of the scientific vessels Nautilus and their ROV pilots for help with sampling and Margot Bligh for help with R-based LC-MS/MS data analysis.

Figure 1D (Created in BioRender. Stuewe, M. (2026) https://BioRender.com/2txphyz).

ChatGPT was used to revise text passages of this manuscript and for writing code for data analysis in R.

## Author contributions

Malin Stüwe (Conceptualization, Data curation, Formal analysis of MALDI-MSI, 16S rRNA FISH and LC-MS/MS data, Visualization, Writing-original draft, Writing-review & editing), Benedikt Geier (Investigation MALDI-MSI and 16S rRNA FISH, Methodology, Writing-review & editing), Dolma Michellod (Investigation LC-MS/MS, Methodology, Writing-review & editing), Miguel Ángel González Porras (Formal analysis of transcriptomics data, Writing-review & editing), Nicole Dubilier (Resources, Supervision, Writing-review & editing), Manuel Liebeke (Conceptualization, Resources, Supervision, Writing-original draft, Writing-review & editing).

## Data availability

MALDI-MSI datasets can be accessed via the METASPACE platform: https://metaspace2020.org/project/bathy_nix_ms. Fluorescence microscopy files, LC-MS/MS datasets and transcriptomics analyses are available on zenodo (https://doi.org/10.5281/zenodo.20200215).

